# Mitochondrial energy dysfunction induces remodeling of the cardiac mitochondrial protein acylome

**DOI:** 10.1101/2021.01.31.429057

**Authors:** Jessica N. Peoples, Nasab Ghazal, Duc M. Duong, Katherine R. Hardin, Nicholas T. Seyfried, Victor Faundez, Jennifer Q. Kwong

## Abstract

Mitochondria are increasingly recognized as signaling organelles because, under conditions of stress, mitochondria can trigger various signaling pathways to coordinate the cell’s response. The specific pathway(s) engaged by mitochondria in response to defects in mitochondrial energy production in vivo and in high-energy tissues like the heart are not fully understood. Here, we investigated cardiac pathways activated in response to mitochondrial energy dysfunction by studying mice with cardiomyocyte-specific loss of the mitochondrial phosphate carrier (SLC25A3), an established model that develops cardiomyopathy as a result of defective mitochondrial ATP synthesis. In heart tissue from these mice, mitochondrial energy dysfunction induced a striking pattern of acylome remodeling, with significantly increased post-translational acetylation and malonylation. Mass spectrometry-based proteomics further revealed that energy dysfunction-induced remodeling of the acetylome and malonylome preferentially impacts mitochondrial proteins. Acetylation and malonylation modified a highly interconnected interactome of mitochondrial proteins, and both modifications were present on the enzyme isocitrate dehydrogenase 2 (IDH2). Intriguingly, IDH2 activity was enhanced in SLC25A3-deleted mitochondria, and further study of IDH2 sites targeted by both acetylation and malonylation revealed that these modifications can have site-specific and distinct functional effects. Finally, we uncovered a novel crosstalk between the two modifications, whereby mitochondrial energy dysfunction-induced acetylation of sirtuin 5 (SIRT5), inhibited its function. Because SIRT5 is a mitochondrial deacylase with demalonylase activity, this finding suggests that acetylation can modulate the malonylome. Together, our results position acylations as an arm of the mitochondrial response to energy dysfunction and suggest a mechanism by which focal disruption to the mitochondrial energy production machinery can have an expanded impact on global mitochondrial function.

## INTRODUCTION

The mitochondrial oxidative phosphorylation system (OXPHOS) is the major source of fuel that drives cellular functions. The critical dependence on mitochondria as an energy source is especially evident in tissues with high-energy demands such as the heart; defects in the mitochondrial energy production machinery underlie a wide range of primary mitochondrial disorders that present with cardiac disease (1,2), and cardiac diseases like heart failure and myocardial infarction are characterized by mitochondrial dysfunction (3–5). Yet, the mitochondrion-intrinsic mechanisms activated in response to primary defects in cardiac mitochondrial energy production are not clear.

Several pathways can communicate mitochondrial dysfunction or stress to the rest of the cell (6), and therefore are candidates for signaling cardiac mitochondrial energy stress. These include the canonical AMP kinase (AMPK) (7,8), reactive oxygen species (ROS) (9), and the mitochondrial unfolded protein response (10) signaling pathways. More recently, metabolites derived from mitochondrial metabolism have also emerged as important mediators of mitochondrial communication (11).

An intriguing group of candidate metabolites is the acyl-coenzymes A’s (acyl-CoAs). These reactive molecules synthesized from metabolic intermediates serve as substrates for acylations, a class of protein post-translational modifications (PTMs) that regulate proteins in pathways ranging from chromatin accessibility to metabolic protein function (12–16). Acylations are reversible covalent additions of acyl groups from reactive acyl-CoAs to the ɛ-amino group of target lysines (13). Acetylation, the addition of the acetyl group of acetyl-CoA to lysine residues, is among the best-studied acylations and modifies a diverse array of proteins across cellular compartments (17–19). In recent years, other acylations have been discovered; these include malonylation, succinylation, and glutarylation (derived from malonyl-CoA, succinyl-CoA, and glutaryl-CoA respectively). These acylations also harbor the potential to modify histone and mitochondrial proteins (20–23), suggesting the tantalizing potential for this class of PTMs to coordinate cellular metabolic state with gene expression and metabolic regulation (12).

While acetylation, malonylation, succinylation, and glutarylation are unified by their status as acylations, individually they may exert different biochemical effects. At physiological pH, acetylation neutralizes the positive charge of modified lysine residues, while malonylation, succinylation, and glutarylation impart a negative charge (23). Additionally, these modifications differ in size and structure (23). As such, acylations may differentially affect protein structure, electrostatic interactions within a protein, or even protein-protein interactions with binding partners. Thus, understanding acylation type- and site-specific effects may be crucial for uncovering the functional consequences of acylations on target proteins.

Interestingly, mitochondrial proteins are particular targets for some acylations. Mass spectrometry studies on mice in which sirtuin 3 (SIRT3, a mitochondrial deacetylase), and sirtuin 5 (SIRT5, a mitochondrial deacylase with demalonylase, desuccinylase, and deglutarylase activity) are knocked out have revealed mitochondrial metabolic pathways as hotspots for these modifications (24,25). However, the causes and functional consequences of acylations and their contributions to disease pathology are open areas of active interest and debate (26–32).

In this study, we sought to identify the mitochondrial stress response pathways specifically engaged by impaired mitochondrial ATP synthesis in the heart. We leveraged our previously described mouse model bearing a temporally-regulated and cardiomyocyte-specific deletion of the mitochondrial phosphate carrier (SLC25A3; *Slc25a3*^*fl/flxMCM*^ mice) (33) as a model of mitochondrial energy dysfunction. SLC25A3 is a major transporter of inorganic phosphate (Pi) into the mitochondrial matrix and an essential component of the mitochondrial ATP synthasome (34,35). SLC25A3 deletion in adult cardiomyocytes causes reduced mitochondrial ATP synthesis and the development of mitochondrial cardiomyopathy like that observed in people with phosphate carrier deficiency (33,36,37). Here, we report that SLC25A3 deletion-induced mitochondrial energy dysfunction drives cardiac acylome remodeling, revealing a novel pathway by which disruption to the mitochondrial energy production machinery may impose control on an expanded network of mitochondrial proteins.

## RESULTS

### Canonical pathways of mitochondrial energy stress signaling are not engaged in response to SLC25A3 deletion in the heart

In the *Slc25a3*^*fl/flxMCM*^ model, *Slc25a3*^*fl/fl*^ mice harboring a loxP targeted Slc25a3 locus were crossed to animals expressing a Cre recombinase under the control of the tamoxifen-inducible and cardiomyocyte-specific α-myosin heavy chain promoter (MCM) (33). To induce SLC25A3 deletion in the adult heart, tamoxifen was administered to 8-week-old adult *Slc25a3*^*fl/flxMCM*^ mice with *Slc25a3*^*fl/fl*^ or MCM animals serving as controls. SLC25A3 deletion recapitulated the previously reported phenotype with impaired cardiac mitochondrial ATP synthesis, sustained tissue ATP content, and subsequent severe cardiac hypertrophy (33).

Using *Slc25a3*^*fl/flxMCM*^ mice, we investigated the stress response pathways activated during mitochondrial energy impairment. We first examined activation of AMPK, an energy-sensing kinase that is activated via phosphorylation at threonine 172 (Thr172) in response to cellular energy stress (38). We performed western blot analyses for cardiac pAMPKα Thr172 in *Slc25a3*^*fl//flxMCM*^ versus *Slc25a3*^*fl/fl*^ control animals over the course of SLC25A3 deletion-induced cardiomyopathy. Total protein lysates were from prepared from hearts collected at 2 (impaired ATP synthesis), 6 (development of hypertrophy), and 10 (severe cardiomyopathy) weeks following tamoxifen induction. No differences were detected in the extent of AMPKα phosphorylation as compared to total AMPK expression levels (Fig. 1A-C), suggesting that AMPK signaling is not a major pathway activated by SLC25A3-deletion induced impairment in mitochondrial ATP synthesis.

**Figure 1.**
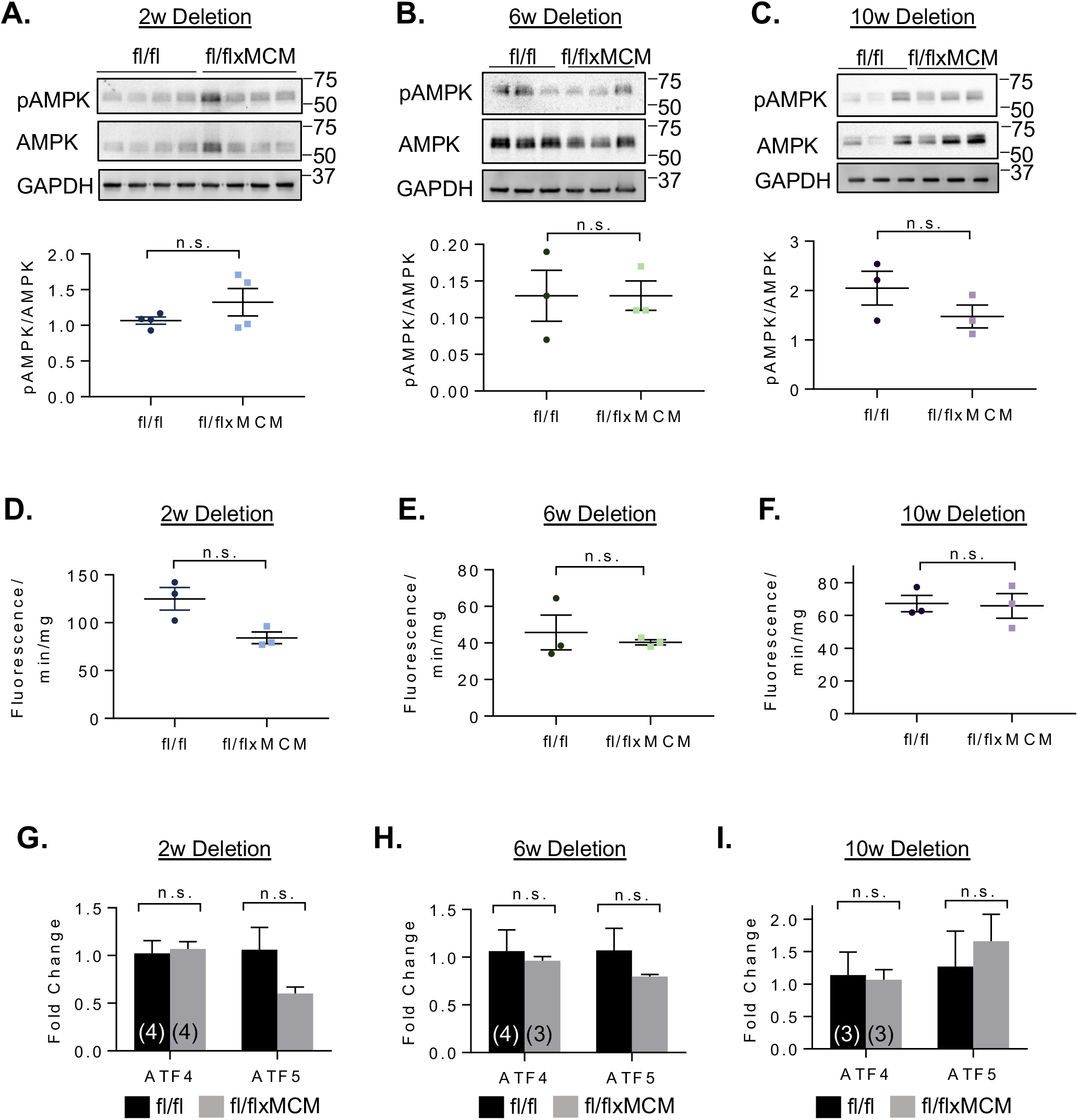
Canonical pathways of mitochondrial energy stress signaling are not activated in response to SLC25A3 deletion in the heart. **A-C.** Western blot analyses and quantification of phosphorylated AMPKα (pAMPK) and total AMPKα in hearts collected from *Slc25a3*^*fl/flxMCM*^ and *Slc25a3*^*fl/fl*^ control mice at 2, 6, and 10 weeks following tamoxifen dosing. GAPDH was used as a protein loading control. **D-F.** Hydrogen peroxide production as measured by an amplex red assay on cardiac mitochondria isolated from *Slc25a3*^*fl/flxMCM*^ and *Slc25a3*^*fl/fl*^ mice at 2, 6, and 10 weeks following tamoxifen dosing. **G-I.** RT-PCR analyses of ATF4 and ATF5 transcript levels in *Slc25a3*^*fl/flxMCM*^ and *Slc25a3*^*fl/fl*^ hearts at 2, 6, and 10 weeks following tamoxifen dosing. Values presented as mean ± SEM. Student’s t test was used for statistical analysis. P<0.05 were considered significant. P<0.05 (*).

Similarly, mitochondrial ROS production is often elevated with mitochondrial dysfunction, and ROS have important signaling functions in the cell (6,9). Using mitochondrial Amplex Red ROS production assays, we measured the rate of hydrogen peroxide synthesis from functional cardiac mitochondria isolated from *Slc25a3*^*fl//flxMCM*^ and *Slc25a3*^*fl/fl*^ mice 2, 6, and 10 weeks following tamoxifen dosing (Fig. 1D-F). Hydrogen peroxide production was unchanged between the two groups (Fig. 1D-F). The absence of enhanced mitochondrial ROS production following SLC25A3 aligns with our previous findings (33), wherein SLC25A3 deletion reduces mitochondrial ATP synthesis, but does not impair respiratory chain function—a major source of mitochondrial ROS.

Finally, we examined the effects of SLC25A3 deletion on the mitochondrial unfolded protein response. This signaling pathway is implicated in mitochondria-to-nucleus retrograde communication and engaged in response to mitochondrial proteotoxic stress and mitochondrial dysfunction (10). Central pathway members are transcription factors ATF4 and ATF5. *Slc25a3*^*fl/flxMCM*^ and *Slc25a3*^*fl/fl*^ hearts displayed similar ATF4 and ATF5 transcript levels (Fig, 1G-I). Taken together, our data suggest that canonical modes of mitochondrial stress signaling are not major contributors to the stress response during SLC25A3 deletion-induced mitochondrial energy defects in the heart. Thus, other response pathways must be activated in this context.

### SLC25A3 deletion induces a unique signature of cardiac protein hyperacylation

Among intermediate metabolites produced via mitochondrial metabolic processes and involved in mitochondrial signaling (11), acyl-CoAs are key substrates for acylations, a class of PTMs that regulate proteins in a variety of pathways and are linked to metabolic dysregulation (12–16). SLC25A3 deletion impairs mitochondrial energy production; if this alters acyl-CoA production, it could impact protein PTM by acylations. Therefore, we assessed the effect of SLC25A3 loss on the cardiac acylome—specifically, acetylation, malonylation, succinylation, and glutarylation. We performed western blot analyses of acetylated-, malonylated-, succinylated-, and glutarylated lysine residues in total protein extracts from *Slc25a3*^*fl/flxMCM*^ versus *Slc25a3*^*fl/fl*^ hearts (Fig. 2). Hearts lacking SLC25A3 had significantly increased levels of acetylation (Fig. 2A) and malonylation (Fig. 2B); succinylation (Fig. 2C) and glutarylation (Fig. 2D) levels were unchanged. These data suggest that SLC25A3 deletion induces remodeling of the cardiac acylome with targeted alterations to the acetylome and malonylome.

**Figure 2.**
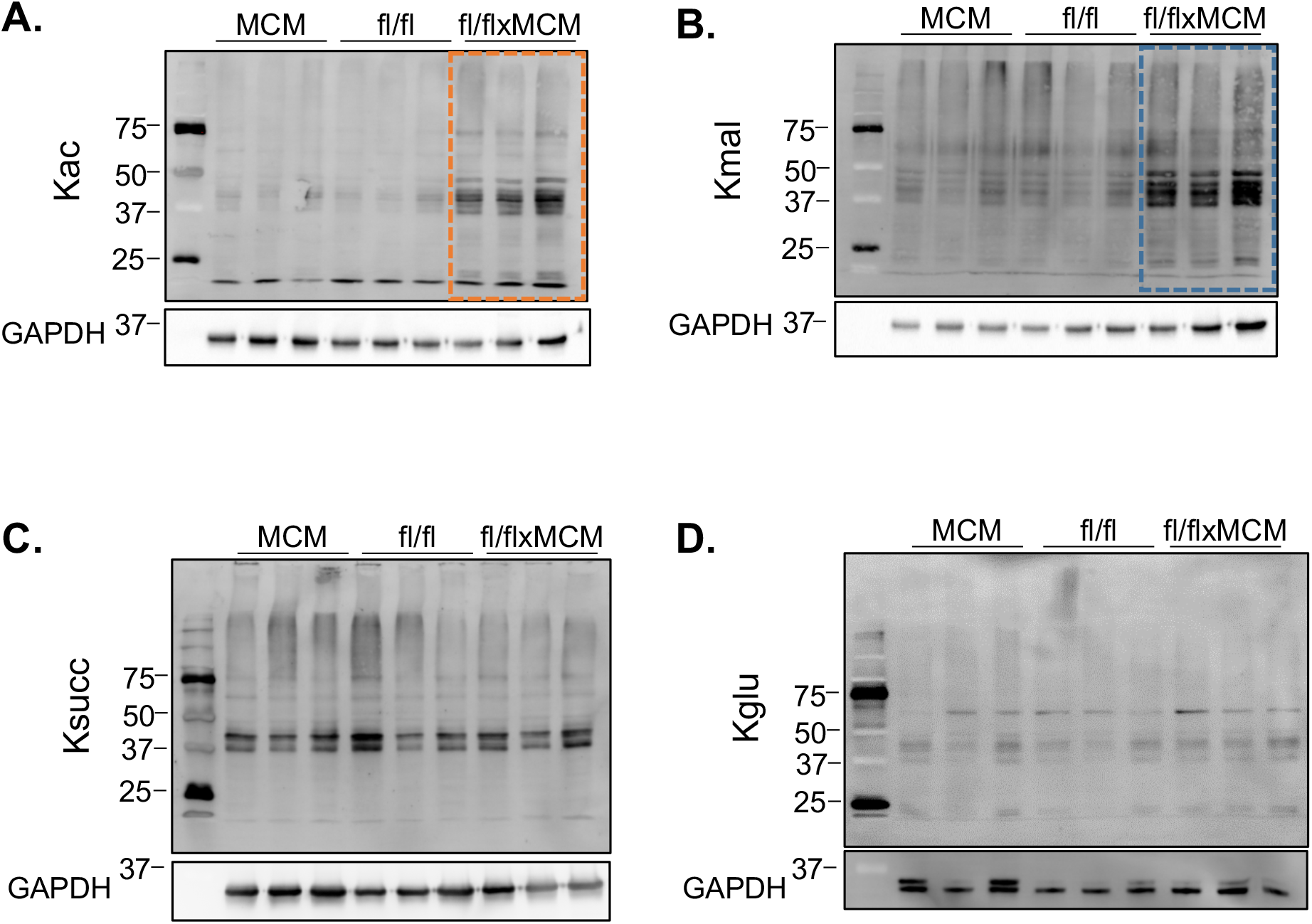
Cardiac SLC25A3 deletion induces a signature of elevated acetylation and malonylation. Western blot analyses of (A) acetylated (Kac), (B) malonylated (Kmal), (C) succinylated (Ksuc), and (D) glutarylated (Kgul) lysines in total protein lysates prepared from *Slc25a3*^*fl/flxMCM*^ and *Slc25a3*^*fl/fl*^ hearts 10 weeks post tamoxifen administration.

### Mitochondrial energy dysfunction-induced acylations preferentially target mitochondrial proteins

We next applied a proteomics approach to identify proteins whose acylation pattern and levels were modified in response to SLC25A3 deletion. Protein extracts from hearts collected 10 weeks after tamoxifen administration (*Slc25a3*^*fl/fl*^ and *Slc25a3*^*fl/flxMCM*^ mice) were immunoaffinity-purified using antibody-conjugated beads to detect acetylated or malonylated lysines. Enriched peptides were analyzed by liquid chromatography tandem mass spectrometry (LC-MS/MS). Differential acylome modifications fell into one of three major categories in response to SLC25A3 deletion: 1) increased acetylation (Fig. 3A; red symbols), 2) decreased acetylation (Fig. 3A; blue symbols), and 3) increased malonylation (Fig. 3B; red symbols). Categories 1 and 2 encompassed a total of 543 peptides representing 208 unique proteins that displayed altered acetylation status in response to SLC25A3 deletion (Fig. 3A and Supplementary Data 1). Within this set of differentially acetylated proteins, 94 proteins harbored lysines that were hyperacetylated in response to SLC25A3 deletion, while 153 proteins harbored lysines that were hypoacetylated (Fig. 3C; Supplemental Data 1). Interestingly, a subgroup of 39 differentially acetylated proteins reflected proteins that had some lysine sites displaying increased acetylation following SLC25A3 deletion, while other sites within the same protein displayed reduced acetylation [Fig. 3C (17+22 intersects)]. Category 3 encompassed 200 differentially malonylated peptides representing 68 proteins, all of which were hypermalonylated after SLC25A3 deletion (Fig. 3B, red symbols; Supplemental Data 2). Additionally, for proteins differentially acylated in response to SLC25A3 deletion, 44 were solely hyperacetylated (category 1), 95 were solely hypoacetylated (category 2), 16 were solely hypermalonylated (category 3). Yet, 22 proteins crossed all three categories (Fig. 3C). These findings reveal a striking and unforeseen complexity of mitochondrial energy dysfunction-induced acylome remodeling.

**Figure 3.**
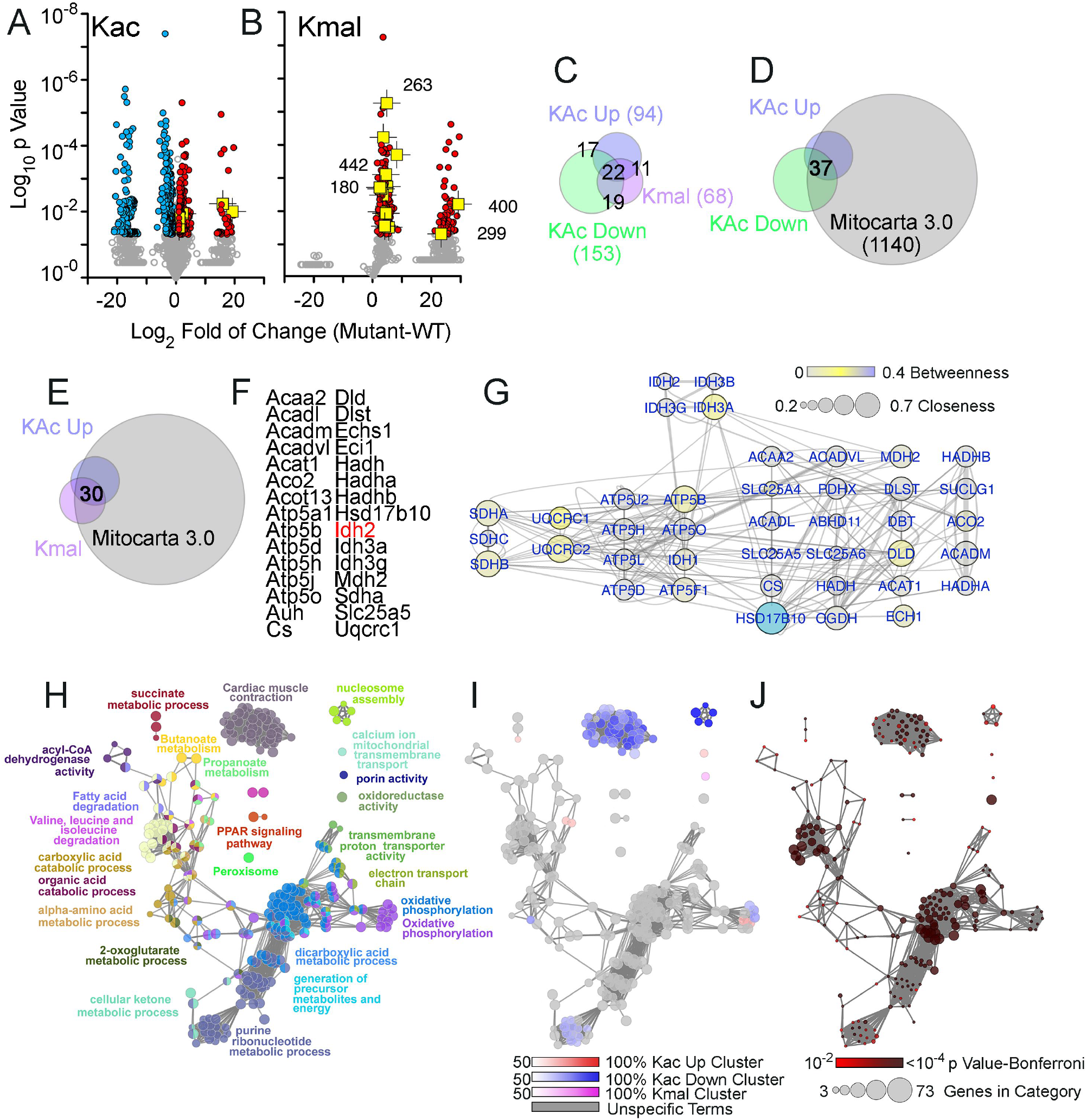
SLC25A3 deletion-induced acylome remodeling preferentially targets mitochondrial ontologies. (A-B) Volcano plots of acetylated proteins (Kac) and malonylated proteins (Kmal). Colored symbols depict peptide hits with a two-fold increase (red symbols) or decrease (blue symbols) and with a p-value <0.05; n=5. IDH2 acylated peptides are depicted in yellow with the lysine modified numbered in B. (C-E) Venn diagrams identifying different categories of acylation and their intersection with the curated Mitocarta 3.0 dataset. (F) List of proteins that exhibited increases in both acylation and malonylation. (G) Interactome of the proteins listed in G. Node connectivity was quantified by betweenness and closeness centrality indexes using Cytoscape. (H-J) ClueGo analysis of the three categories of acylome modifications: increased acetylation, decreased acetylation, and increased malonylation. The GO Biological process and KEGG databases were queried with a threshold Bonferroni-corrected p value below 0.01. H depicts all proteins in the three categories analyzed simultaneously. I presents the percent contribution of each of the three acylation categories to the ontology term presented in H. J depicts the p value and number of genes per ontology term presented in H.

To determine the subcellular distribution of proteins targeted by mitochondrial energy dysfunction-induced acylations, we analyzed our SLC25A3 deletion-induced acetylated and malonylated proteins according to their annotation in the Mitocarta3.0 database (39). Most hyperacetylated proteins were annotated to the mitochondrion (Fig. 3D, 91%), as were most hypermalonylated proteins (Fig. 3E, 73%). This preferential modification of mitochondrial proteins by energy dysfunction-induced acetylation and malonylation was confirmed by western blot analyses for acetylated and malonylated lysine residues on mitochondrial protein extracts from *Slc25a3*^*fl/flxMCM*^ versus *Slc25a3*^*fl/fl*^ hearts (Supplemental Fig. 1). In contrast, a reduced percentage of hypoacetylated proteins were annotated to the Mitocarta 3.0 database (46%). Collectively, these results indicate that SLC25A3 deletion-induced remodeling of the cardiac acetylome and malonylome occurs predominantly in the mitochondrial compartment.

We focused on 30 proteins with increased acetylation and malonylation in mutant hearts. All of these proteins were annotated to mitochondria (Fig. 3E and F). These 30 proteins belong to a high-connectivity interactome defined by protein-protein interactions as well as functional interactions annotated in the REACTOME database (40) (Fig. 3G). This interactome includes subunits of the respiratory chain complexes II, III, and V; all three ADP-ATP translocators (SLC25A4-6); enzymes involved in mitochondrial fatty acid oxidation such as HADHA and HADHB; and enzymes of the Krebs cycle such as isocitrate dehydrogenase 2 (IDH2). IDH2 is of particular interest as a critical regulatory step in the Krebs cycle and for being an example of a protein highly acetylated and malonylated in response to SLC25A3 deletion (Fig. 3A-B, yellow symbols).

Finally, we asked whether the three acylation categories segregated into distinct molecular mechanisms. We used the ClueGo bioinformatic tool to assess gene ontologies using the three categories as gene sets. The most significant terms identified across categories were carboxylic acid metabolic process (GO:0019752, Bonferroni-corrected p = 3.7E-41) and the KEGG term oxidative phosphorylation (KEGG:00190, Bonferroni-corrected p = 4.3E-34; Fig. 3H-I, gray colors and Supplemental Data 3). However, proteins with reduced acetylation in response to SLC25A3 deletion were uniquely enriched in the DNA packaging annotated term (GO:0006323, Bonferroni-corrected p = 0.0057) and cardiac muscle contraction (GO:0060048, Bonferroni-corrected p = 0.000004; Fig. 3H-J). Malonylated proteins were enriched in the annotated term porin activity, represented by the genes *Vdac1*, *Vdac2*, and *Vdac3* (GO: 0015288, Bonferroni-corrected p = 0.001). These results demonstrate that proteins newly acylated upon mitochondrial energy dysfunction center around mitochondrial annotated ontologies, while proteins losing acylations are distinguished by non-mitochondrial terms such as DNA packaging and muscle contraction.

### Mitochondrial acetylation and malonylation have site-specific and functionally distinct effects

We next explored the functional effects of a subset of proteins targeted for acylome remodeling. As described above, IDH2 is both hyperacetylated (5 sites) and hypermalonylated (9 sites) in *Slc25a3*^*fl/flxMCM*^ hearts compared to *Slc25a3*^*fl/fl*^ controls (Fig. 4A-B). We validated the mass spectrometry findings via immunoprecipitation (IP) of IDH2 from heart protein extracts and immunoblotting with antibodies against acetylated or malonylated lysines (Fig. 4C).

**Figure 4.**
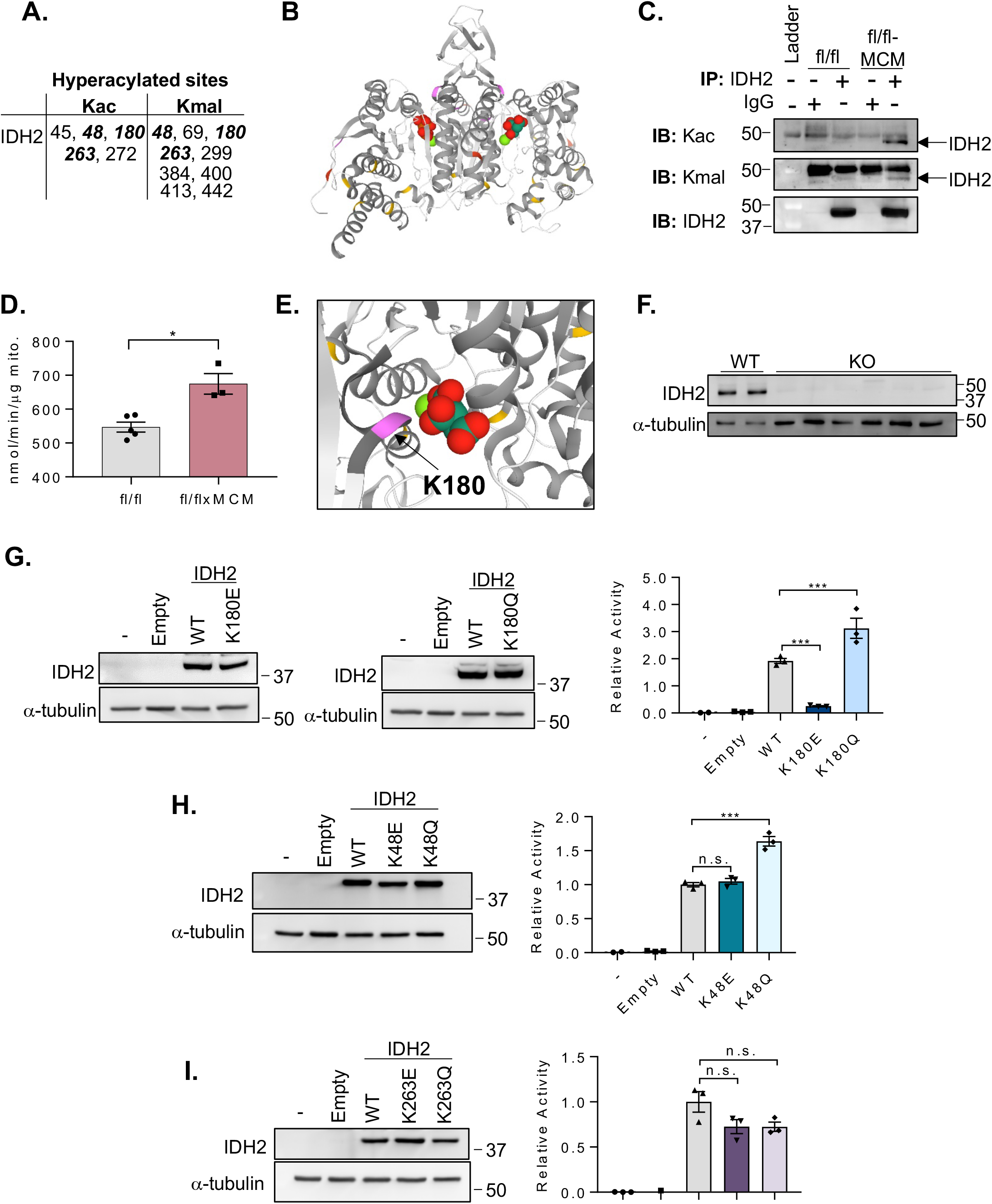
Site-specific expression and activity of IDH2 acetylation and malonylation mimics. (A) IDH2 lysine residues hyperacetylated (Kac) and hypermalonylated (Kmal) in response to SLC25A3 deletion. Residues modified by both PTMs in bold. (B) Structure of the IDH2 dimer bound to isocitrate. Lysine resides acetylated (red), malonylated (yellow), or targeted by both PTMs (purple) in response to SLC25A3 deletion are indicated. (C) IDH2 was immunoprecipitated from protein lysates from *Slc25a3*^*fl/fl*^ or *Slc25a3*^*fl/flxMCM*^ hearts and probed by immunoblotting by anti-Kac or anti-Kmal antibodies to confirm enhanced acetylation/malonylation of IDH2 in response to SLC25A3 deletion. Immunoblotting with anti-IDH2 antibodies was conducted to validate pull down of IDH2. Immunoprecipitation was also conducted with IgG antibodies as a control. (D) IDH2 activity assay performed on cardiac mitochondria isolated from *Slc25a3*^*fl/fl*^ and *Slc25a3*^*fl/flxMCM*^ animals 10 weeks post-tamoxifen administration. (E) Representation of K180 (purple) and isocitrate in the IDH2 binding pocket. (F) Western blot analyses of IDH2 expression in WT and IDH2 KO clones. Α-tubulin was used as a protein loading control. (G) Western blot analyses of IDH2 expression in IDH2 KO cells re-expressing either the pcDNA3.1 vector alone (empty), WT IDH2, IDH2 K180E, or IDH2 K180Q constructs, and the IDH2 activity assays on mitochondria isolated from these cell lines. (H) Western blot analyses of IDH2 expression in IDH2 KO cells re-expressing either the empty vector, WT IDH2, IDH2 K48E, or IDH2 K48Q constructs, and the IDH2 activity assays on mitochondria isolated from these cell lines. (I) Western blot analyses of IDH2 expression in IDH2 KO cells re-expressing either the empty vector, WT IDH2, IDH2 K263E, or IDH2 K263Q constructs, and the IDH2 activity assays on mitochondria isolated from these cell lines. For all graphs, values presented as mean ± SEM. For (D-F), one way ANOVA followed by Welch’s test was used for statistical analysis with P<0.05 considered significant. For (G), a Student’s t test was used for statistical analysis. P<0.05 were considered significant. P<0.05 (*), and P<0.001 (***).

Acylations of mitochondrial metabolic enzymes, including IDH2, are reported to be inhibitory (41–43). We therefore hypothesized that hyperacylations of IDH2 would suppress its function. To test this, we examined IDH2 expression and activity in mitochondria isolated from *Slc25a3*^*fl/flxMCM*^ versus *Slc25a3*^*fl/fl*^ control hearts. While SLC25A3 deletion had no effect on overall cardiac mitochondrial IDH2 protein levels (Supplemental Fig. 2), surprisingly, mitochondria isolated from *Slc25a3*^*fl/flxMCM*^ hearts had significantly elevated IDH2 activity as compared to mitochondria from *Slc25a3*^*fl/fl*^ controls (Fig. 4D).

This striking finding prompted us to examine the structural positioning of the hyperacylated lysines (Fig. 4B). We determined that K180, a lysine targeted by both acetylation and malonylation, lies in close proximity to the isocitrate substrate binding pocket (Fig. 4E), suggesting that modification at this site may modulate enzyme function. Two other sites, K48 and K263, were also hyperacetylated and hypermalonylated in response to SLC25A3 deletion. We leveraged these three modification targets to determine whether functional effects of acylations are site-specific. We generated mutant IDH2 constructs simulating constitutive acetylation (mutation of lysine to glutamine; Q) (44), or constitutive malonylation (mutation of the target lysine to glutamic acid; E) (25). Wild type (WT) IDH2 or the acylation mutant constructs were transiently re-expressed into an HEK293 IDH2 knockout (KO) cell line generated by CRISPR-Cas9 mediated gene editing (Fig. 4F), which were then assayed for IDH2 activity on mitochondria. IDH2 KO mitochondria, as well as mitochondria from KO cells expressing the pcDNA3.1 vector alone (empty), displayed minimal IDH2 activity; in contrast, mitochondria isolated from KO cells re-expressing WT IDH2 displayed significantly elevated IDH2 function (Fig. 4G-I). In line with previous reports that K263 acetylation has no impact on IDH2 function (42), mitochondria isolated from KO cells re-expressing the IDH2 K263Q acetylation mimic displayed IDH2 activity levels similar to mitochondria from WT re-expressing cells (Fig. 4I). Mitochondria isolated from KO cells re-expressing the K263E constitutive malonylation mutant also displayed activity levels similar to WT mitochondria (Fig. 4I). These findings collectively suggest that acetylation and malonylation at K263 does not impact IDH2 function.

For modification at K48, mitochondria from KO cells re-expressing the IDH2 K48E mutant displayed similar IDH2 activity levels as WT IDH2 re-expressing mitochondria, suggesting that malonylation at this site has no functional effects (Fig. 4H). Surprisingly, however, mitochondria from KO cells expressing the constitutive acetylation K48Q mutant displayed elevated IDH2 activity levels as compared to WT re-expressing mitochondria (Fig. 4H), suggesting that acetylation at this site enhances function. For SLC25A3-deletion responsive acylations at K180, mitochondria re-expressing the constitutive malonylation K180E mutant displayed significantly reduced IDH2 activity, while re-expression of the constitutive acetylation K180Q mutant at this resulted in elevated activity (Fig. 4G), suggesting that acetylation and malonylation at K180 have opposing effects.

Our data suggest that, cumulatively, SLC25A3 deletion-induced hyperacetylation and hypermalonylation of IDH2 enhances enzyme activity, and that acetylation and malonylation at individual sites may have functionally distinct effects.

### Mitochondrial malonylation is modulated by acetylation

Our analyses of the pathways engaged in response to selective ablation of SLC25A3 in the heart revealed that defective mitochondrial ATP synthesis causes specific hyperacetylation and hypermalonylation of mitochondrial proteins (Fig. 3). While alterations in mitochondrial protein acetylation are associated with cardiac mitochondrial energy dysfunction and various models of cardiac disease (19,26,41), our discovery that mitochondrial ATP synthesis deficiency induces a concomitant increase in mitochondrial protein malonylation was striking and unanticipated.

To characterize the mechanisms underlying energy dysfunction-induced malonylation, we first examined the impact of SLC25A3 deletion on pathways (modification addition and removal) that modulate mitochondrial protein malonylation status. Malonylation of mitochondrial proteins is thought to occur by acyltransferase-independent chemical reactions, facilitated by permissive conditions within the mitochondrial matrix (45,46). Chemical malonylation may therefore be modulated by malonyl-CoA substrate availability. To test this, we applied targeted liquid-chromatography mass spectrometry for malonyl-CoA to *Slc25a3*^*f/flxMCM*^ and *Slc25a3*^*fl/fl*^ control hearts. Remarkably, SLC25A3 deletion promoted significantly elevated malonyl-CoA levels in the heart (Fig. 5A). Because the mitochondrial malonylome is also regulated by the SIRT5, a mitochondrial demalonylase, (45), we examined the effects of SLC25A3 deletion on SIRT5. Total levels of SIRT5 were unaltered by SLC25A3 deletion (Fig. 5B). However, further examination of our mass spectrometry data revealed that SIRT5 is hyperacetylated at K203 in response to SLC25A3 deletion. This finding was confirmed by IP and immunoblotting (Fig. 5C).

**Figure 5.**
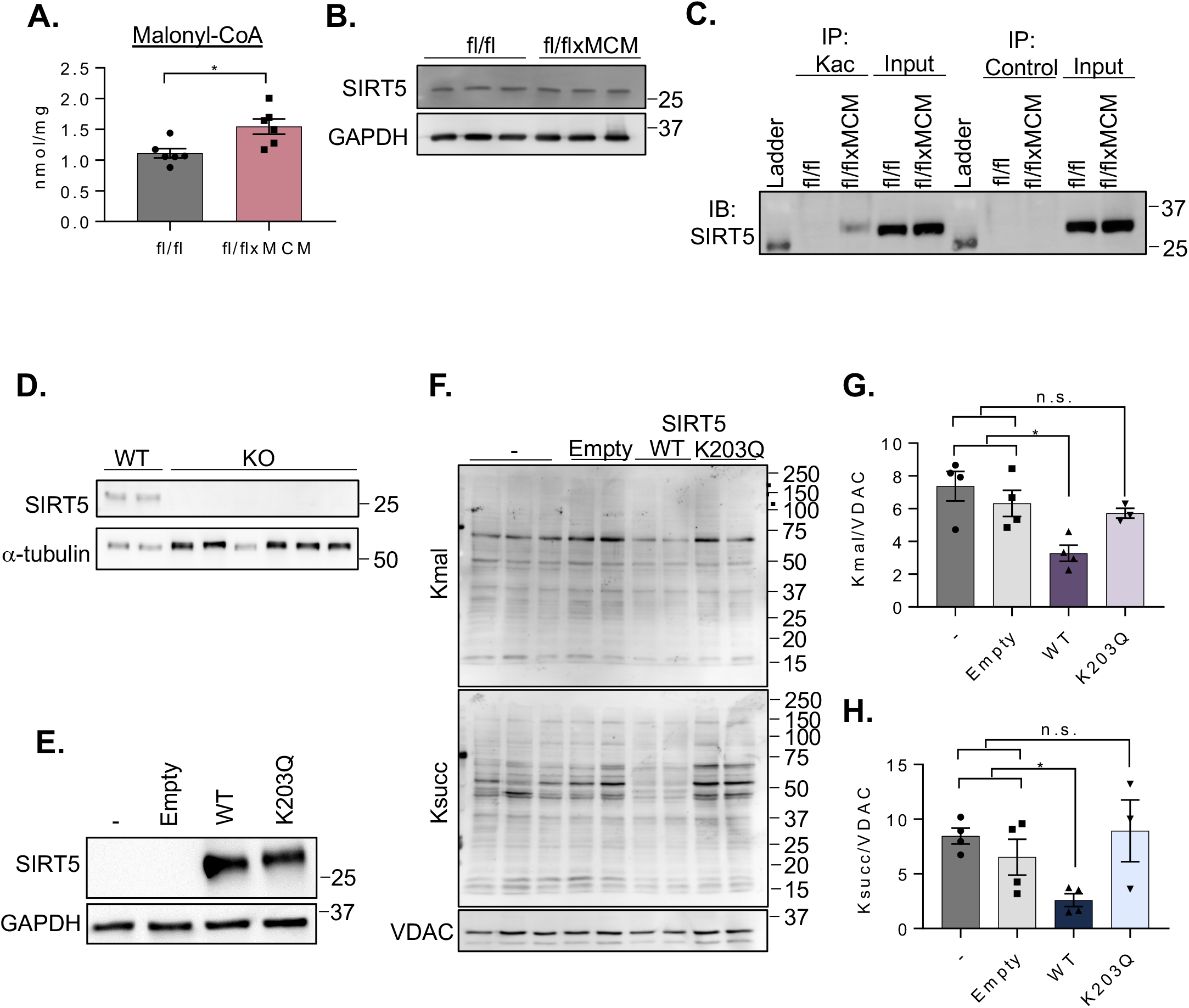
Acetylation of SIRT5 modulates malonylation status of mitochondrial proteins. (A) Cardiac malonyl-CoA levels in *Slc25a3*^*fl/fl*^ versus *Slc25a3*^*fl/flxMCM*^ mice at 10 weeks post-tamoxifen administration. (B) Western blot analysis of SIRT5 expression in total heart protein lysates from *Slc25a3*^*fl/fl*^ versus *Slc25a3*^*fl/flxMCM*^ animals. GAPDH was used as a loading control. (C) Immunoprecipitation confirmation of enhanced SIRT5 acetylation in *Slc25a3*^*fl/flxMCM*^ versus *Slc25a3*^*fl/fl*^ hearts. Acetylated proteins were immunoprecipated with anti-Kac antibodies followed by immunoblotting with an antibody against SIRT5. An anti-HA antibody was used as an immunoprecipitation control. (D) Western blot analysis of SIRT5 expression in HEK293 WT versus SIRT5 KO clones. α-tubulin was used as a protein loading control. (E) Western blot analysis of SIRT5 expression in SIRT5 KO cells re-expressing a pcDNA3.1 empty vector, WT SIRT5 or, the SIRT5 K203Q mutant. GAPDH was used as a loading control. (F) Representative western blot of protein lysine malonylation (Kmal) and succinylation (Ksuc) of mitochondria isolated from SIRT5 KO cells, or SIRT5 KO cells re-expressing the empty vector, WT SIRT5, or SIRT5 K203Q. VDAC was used as a loading control. (G) Quantification of protein malonylation in SIRT5 KO cells re-expressing the WT and K203Q SIRT5 constructs. (H) Quantification of protein succinylation in SIRT5 KO cells re-expressing the WT and K203Q SIRT5 constructs. One way ANOVA followed by Welch’s test was used for statistical analysis with P<0.05 considered significant. P<0.05 (*).

To investigate the functional impact of SIRT5 K203 acetylation, we generated SIRT5 KO HEK293 cell lines by CRIPSR/Cas9-mediated gene deletion (Fig. 5D) and re-expressed either WT SIRT5 (pcDNA3.1-SIRT5-WT), or a SIRT5 K203Q mutant (pcDNA3.1-SIRT5-K203Q) to mimic enforced lysine acetylation (Fig. 5E). We then examined the effects of these constructs on global protein malonylation. As anticipated, expression of a pcDNA3.1 empty vector control in SIRT5 KO cells had no effect on malonylation levels, while overexpression of WT SIRT5 reduced the degree of malonylation observed in total cell lysates (Fig. 5F-H). Surprisingly, overexpression of the SIRT5 K203Q enforced-acetylation mutant significantly depressed the extent of malonylation observed in SIRT5 KO cells (Fig. 5F-G), suggesting that acetylation at K203 inhibits SIRT5 function. Because SIRT5 also possesses desuccinylase activity, we examined the effects of WT SIRT5 and the K203Q enforced acetylation mutant on protein succinylation. Cells overexpressing WT SIRT5 displayed reduced total succinylation as compared to untransfected and empty-vector controls, but overexpression of the K203Q mutant did not alter total protein succinylation (Figs 5F, H). Taken together, these data suggest that acetylation of SIRT5 at K203 impairs the ability of SIRT5 to function as a deacylase.

## DISCUSSION

Acylations like acetylation and malonylation are well-documented to target mitochondrial proteins (41,47,48) and are linked to metabolism. In fact, metabolic pathways like the tricarboxylic acid cycle and β-oxidation are highly acylated when mitochondrial deacylation is disrupted, suggesting that acylations may play a critical role in modulating mitochondrial metabolic function (23–25,49). Here, we found that the converse can also occur—that primary defects in mitochondrial energetics (via induction of impaired mitochondrial ATP production through SLC25A3 deletion) can dictate specific changes to the cardiac acylome.

Our results directly demonstrate that, in vivo and in the context of the heart, mitochondrial energy dysfunction caused by impaired mitochondrial ATP synthesis does not engage canonical modes of mitochondrial stress signaling such as the AMPK, ROS, or mitochondrial unfolded protein response pathways. Instead, we found that impaired mitochondrial ATP synthesis induces cardiac acylome remodeling with a specific signature of elevated mitochondrial acetylation and malonylation (summarized in Fig. 6). Acylations present an attractive potential link between cellular metabolism and gene regulation (16). Acetylation, for example, can modify histone proteins to regulate gene expression by modulating chromatin accessibility (17,20-22). Our observations of elevated acetylation and malonylation of cardiac proteins led us to initially postulate that histone proteins could be a major target of mitochondrial energy dysfunction-induced acylations. Surprisingly, however, mass spectrometry analyses revealed that these energy dysfunction-responsive PTMs were concentrated in the mitochondrial compartment, with proteins involved in metabolism (carboxylic acid metabolism and oxidative phosphorylation) representing some of the top modified pathways.

**Figure 6.**
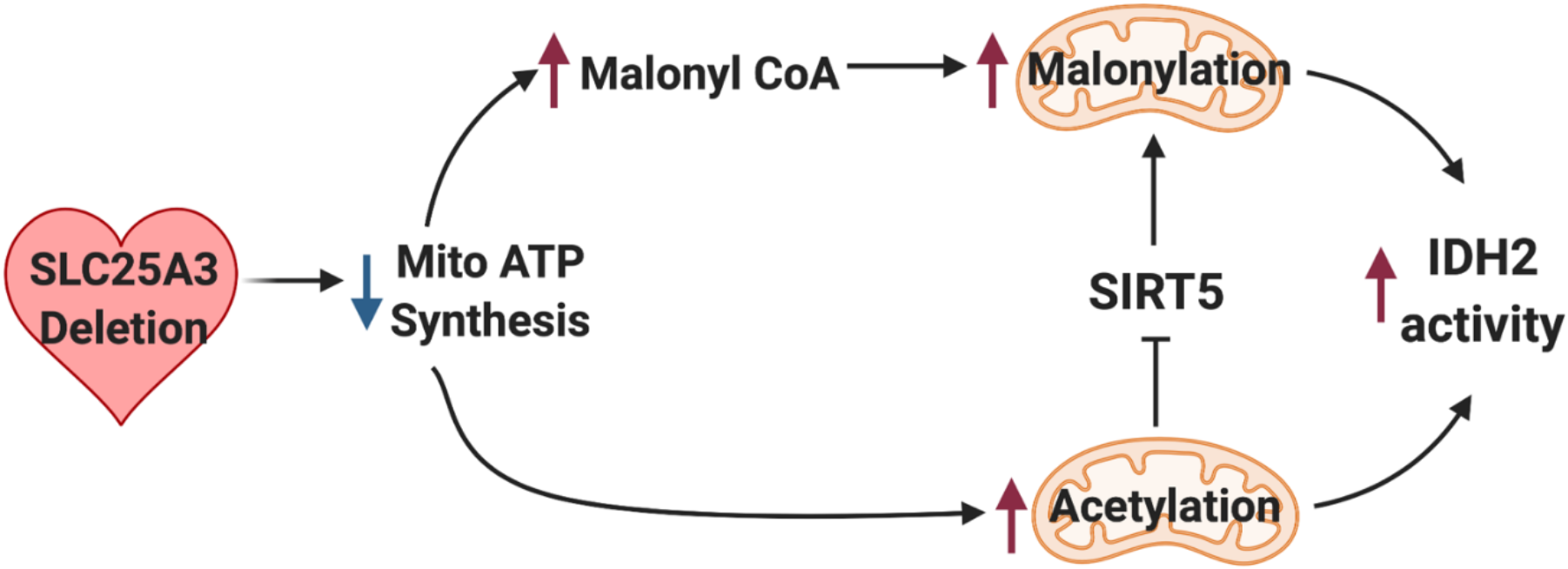
Model of mitochondrial energy dysfunction-induced enhancement of mitochondrial protein acylation. Deletion of SLC25A3 deletion in the heart causes impaired mitochondrial ATP synthesis resulting in mitochondrial energy dysfunction. This leads to mitochondrial protein hyperacetylation and hypermalonylation. Loss of SLC25A3 increases cardiac malonyl-CoA levels which can support enhanced malonylation. Additionally, SLC25A3 deletion results in SIRT5 acetylation which inhibits SIRT5 function as a second mechanism to potentiate mitochondrial malonylation. Finally, mitochondrial energy dysfunction-induced acetylation and malonylation of IDH2 increases IDH2 activity.

While acylations can exert different regulatory consequences (50), in the context of metabolism, acetylation and malonylation of mitochondrial proteins are largely thought to be inhibitory (41). Here, we found that SLC25A3 deletion enhances both acetylation and malonylation of IDH2 and that mitochondrial IDH2 activity is actually increased in the absence of SLC25A3. Intriguingly, we found that the subset of IDH2 lysine residues (K48, K180, and K263) targeted by both PTMs exerted different functional consequences. Mutagenesis studies indicated that IDH2 acylation at some sites can enhance function. Given that acetylation neutralizes the positive charge of target lysines, while malonylation imparts a negative charge and is a structurally larger modification (23), it is not surprising that we found that acetylation and malonylation may elicit differing effects on target proteins—even within the same protein target. At present, we do not know how all of the identified energy dysfunction-induced acylations of IDH2 are integrated to produce the increased activity that we observed in *Slc25a3*^*fl/flxMCM*^ hearts. However, the ultimate functional consequences of these PTMs may depend in part on the type of modification, the specific sites modified, the occupancy rates of each modification, and the cooperative effects of these modifications. Thus, future work must deploy quantitative methods to gain a more complete picture of how energy dysfunction-induced acylations modify target proteins.

Enhanced acetylation occurs in a number of different of cardiac disease models (19,26), as well as in a model of impaired complex I assembly (51). We anticipated that mitochondrial energy dysfunction due to SLC25A3 deletion would induce similar acetylome changes. We were surprised, however, to find that SLC25A3 deletion also resulted in a high degree of mitochondrial protein malonylation. Our data point to two major pathways that contribute to energy dysfunction-induced malonylation (Fig. 6). First, our observation that SLC25A3-deleted hearts display significantly elevated levels of malonyl-CoA suggests that increased malonyl-CoA availability could facilitate chemical additions that lead to the observed increases in malonylation. Second, acetylation of SIRT5 at a novel site (K203) reduced this enzyme’s ability to demalonylate proteins, suggesting that impaired SIRT5 function could also contribute to the observed hypermalonylation. How SIRT5 acetylation inhibits function is unclear, as K203 lies outside of the enzyme’s catalytic domain (52). Structurally, however, SIRT5 is composed of two globular domains, the smaller of which harbors a zinc binding site that is essential for enzyme structure/function and substrate recognition (52,53). K203 resides within this smaller globular domain, and acetylation at this site could impact substrate recognition and zinc binding. Importantly, however, our findings suggest a novel crosstalk between acetylation and malonylation, whereby acetylation can modulate the malonylome through SIRT5 inhibition.

Finally, in addition to the energy dysfunction-induced increases in mitochondrial protein acetylation and malonylation, our proteomics study revealed a surprising and unexpected decrease in acetylation of a small group of non-mitochondrial proteins implicated in DNA packaging and cardiac muscle contraction. At present, the functional impact and mechanisms underlying this energy dysfunction-responsive reduction in acetylation are unknown. However, alterations to these pathways may contribute to the overall cardiac response to mitochondrial energy deficits.

In conclusion, we found that mitochondrial energy dysfunction due to defective mitochondrial ATP synthesis initiates a novel stress response independent of canonical AMPK, ROS, and mitochondrial unfolded protein response signaling. Hyperacylation of mitochondrial proteins has been implicated in a number of disease contexts (19,26,41,54–56), and the expanded network of mitochondrial proteins targeted by mitochondrial energy dysfunction-induced acylations that we uncovered suggests a mechanism whereby focal defects in the mitochondrial ATP synthasome (via SLC25A3 deletion) can control an expanded network of mitochondrial proteins. Notably, recent work using a striated muscle-specific double-knockout model of SIRT3 and carnitine acetyltransferase has called in to question the role of enhanced mitochondrial acetylation, if any, in the transverse aortic constriction surgical model of cardiac pressure overload-induced heart failure (31); thus, future work should determine the functional consequences of hyperacetylation and hypermalonylation in the context of primary mitochondrial dysfunction and cardiac mitochondrial ATP synthesis defects. Understanding if acylations merely serve as post-translational biomarkers/signatures of altered metabolism, or if they have context-specific and disease-specific roles of promoting pathology will critical next steps as we move to a greater understanding of how mitochondria communicate dysfunction and aim to leverage these pathways for future therapies.

## EXPERIMENTAL PROCEDURES

### Animal Models

The *Slc25a3* loxP targeted mice (*Slc25a3*^*fl/fl*^) and the αMHC-MerCreMer animals expressing a tamoxifen inducible Cre recombinase under the control of the cardiomyocyte specific alpha-myosin heavy chain promoter (MCM) were described previously (33,57). *Slc25a3*^*fl/flxMCM*^ mice were generated by crossing *Slc25a3*^*fl/fl*^ mice to the MCM animals as previously described (33). *Slc25a3* deletion was induced in 8-week-old *Slc25a3*^*fl/flxMCM*^ animals by intraperitoneal injections of tamoxifen (25 mg/kg for 5 consecutive days). As controls, 8-week-old *Slc25a3*^*fl/fl*^ and MCM control animals, harboring either the Slc25a3 targeted locus or MCM transgene alone, were subjected to the same tamoxifen dosing regimen. All animal experiments were approved and performed in accordance with Emory University’s Institutional Animal Care and Use Committee.

### Cell Culture, Plasmid Vectors, and Transfection

Human embryonic kidney 293 (HEK293) cell lines were cultured in Dulbeco’s minimal essential media supplemented with 10% bovine growth serum and 1% penicillin/streptomycin antibiotics (100units/mL pencillin and 100ug/mL streptomycin) in 5% CO_2_ atmosphere at 37°C. IDH2 and SIRT5 knockout (KO) HEK293 cell lines were generated by CRIPSR-Cas9 mediated gene deletion (Synthego) and individual KO clonal lines were isolated by dilution cloning.

PcDNA3.1-IDH2 and pcDNA3.1-SIRT5 vectors were generated by custom DNA synthesis (Genewiz). Site directed mutagenesis (Q5 Site Directed Mutagenesis Kit; New England Biolabs) was conducted to generate the IDH2 K48Q, K180Q, and K263Q constitutive acetylation mimics, the IDH2 K48E, K180E, and K263E constitutive malonylation mimics, as well as the SIRT5-K203Q constitutive acetylation mutant.

For the re-expression of IDH2 and SIRT5 constructs into IDH2 and SIRT5 KO cells, a single IDH2 or SIRT5 KO clonal line was selected for re-expression studies. Cells were transiently transfected with IDH2 and SIRT5 vectors using Lipofectamine 3000 (Invitrogen) as described by the manufacturer. Levels of IDH2 or SIRT5 overexpression was assessed by immunoblotting using specific antibodies against IDH2 and SIRT5 as described below.

### Liquid Chromatography Tandem Mass Spectrometry and Analysis

Mouse cardiac proteins were extracted and digested using a protocol adapted from (58). Briefly, tissue was solubilized in urea lysis buffer (8M urea, 10 mM Tris, 100 mM NaH2PO4, pH 8.5) supplemented with protease and phosphatase inhibitors (Thermo Fisher Scientific) using a Bullet Blender (Next Advance) according to the manufacturer’s protocol. Protein homogenates were sonicated and cleared by centrifugation. Subsequently proteins were diluted with 50 mM NH4HCO3 to a final concentration of 2 M urea, reduced with 1 mM 1,4-dithiothreitol for 30 min and alkylated with 5 mM iodoacetamide for 30 min. Protein samples were then digested with 1:100 (w/w) Lys-C (Wako) at room temperature (r.t.) for 4 hours and then further digested overnight with 1:50 (w/w) trypsin (Promega) at r.t. Resulting peptides were desalted with Oasis HLB columns (Waters) and dried under vacuum.

Acylated (either acetylated or malonylated) peptides were enriched by immunoaffinity purification using the PTMScan Acetyl-Lysine or PTMScan Malonyl-Lysine Kits (Cell Signaling Technology) according to the manufacturer’s protocol. Acylated peptides were further purified by desalting using C18 StageTips. Finally, peptides were reconstituted in 0.1% trifluoroacetic acid. The protocol for data acquisition by LC-MS/MS was adapted from (58). Briefly, 2 μl of the reconstituted peptides were separated on a self-packed C18 (1.9 μm Dr. Maisch, Germany) fused silica column (25 cm x 75 μM internal diameter; New Objective, Woburn, MA) by a Dionex Ultimate 3000 RSLCNano. Elution was performed over a 120 minute gradient at a rate of 300 nL/min with buffer B ranging from 3% to 80% (buffer A: 0.1% formic acid in water, buffer B: 0.1 % formic in acetonitrile). Peptides were analyzed on an Orbitrap Fusion Tribrid Mass Spectrometer (ThermoFisher Scientific). The mass spectrometer cycle was programmed to collect at the top speed for 3 second cycles. The MS scans (300-1500 m/z range, 200,000 AGC, 50 ms maximum ion time) were collected at a resolution of 120,000 at m/z 200 in profile mode and the HCD MS/MS spectra (15000 resolution, 1.6 m/z isolation width, 30% collision energy, 10,000 AGC target, 35 ms maximum ion time) were detected in the Orbitrap. Dynamic exclusion was set to exclude previously sequenced precursor ions for 20 seconds within a 10 ppm window. Precursor ions with +1, and +8 or higher charge states were excluded from sequencing.

MS/MS spectra were evaluated using Proteome Discoverer 2.1.1.21. The raw files were search using a Uniprot *Mus musculus* protein database (downloaded April 2015 with 53289 target sequences). Search parameters included fully tryptic specificity, 20 ppm precursor ion tolerance, 0.05 Da product ion tolerance, static modification for carbamidomethyl cysteine (+57.02146) and dynamic modifications for oxidized methionine (+15.99492), deamidated asparagine and glutamine (+0.98402). Additional dynamic modifications of lysine acetylation (+42.01057) and malonylation (+86.00039). All peptide spectral matches were filtered via the percolator node.

Gene ontology analyses were performed with the Cytoscape Cluego plugin (59). *In silico* interactome data were downloaded from Genemania physical interactions and REACTOME curated interactions were included in the analysis (60) and processed in Cytoscape v3.5.1 (61). Interactome connectivity graph parameters were generated in Cytoscape. Physical interactions were curated from the following sources and their strength represented by the edge thickness (62–66)

### LC/MS/MS Quantitation of Malonyl-CoA

Working calibration standard mix was prepared for malonyl CoA. The calibrators and sample were spiked with a mixture of heavy isotope-labeled internal standards for malonyl CoA. The CoAs were then extracted using Oasis HLB 1cc (30 mg) Extraction Cartridges (Waters Corporation). The extracted samples were dried under nitrogen, reconstituted in 10 mM Ammonium carbonate, pH 9.5. and separated on an Acquity UPLC BEH C18 2.1 x 50 mm, 1.7 μm column, (Waters Corporation) using a 2.35 min linear gradient with 10 mM ammonium carbonate, pH 9.5 and ACN as eluents. Quantitation of malonyl-CoA was achieved using multiple reaction monitoring of calibration solutions and study samples on an Agilent 1290 HPLC/6490 triple quadrupole mass spectrometer. The mass spectrometer was operated in positive ion mode using electrospray ionization with an ESI capillary voltage of 3500V. The electron multiplier voltage was set to 400V. The ion transfer tube temperature was 325 °C and vaporizer temperature was 325 °C. The ESI source sheath gas flow was set at 10L/min. The mass spectrometer was operated with a mass resolution of 0.7 Da, and N2 collision gas pressure was 30 psi for the generation and detection of product ions of each metabolite. For data processing, the raw data was processed using Mass hunter quantitative analysis software (Agilent). Calibration curves (R^2^ = 0.99 or greater) were either fitted with a linear or a quadratic curve with a 1/X or 1/X^2^ weighting.

### Mitochondrial Isolation

Cardiac mitochondria used for immunoblotting and immunoprecipitation were isolated by differential centrifugation as previously described (33) using MS-EGTA buffer (225 mM mannitol, 75 mM sucrose, 5 mM HEPES, and 1 mM EGTA, pH 7.4) supplemented with 2.4 mg/ml nicotinamide, Trichlostatin A, and a combined phosphatase and protease inhibitor cocktail (Thermo Fisher).

### Electrophoresis and Immunoblotting

For western blot analyses, total protein extracts were prepared from hearts homogenized and solubilized in radioimmunoprecipitation assay (RIPA) buffer supplemented with 2.4 mg/ml nicotinamide and a combined protease and phosphatase inhibitor cocktail (Thermo Fisher). Cardiac mitochondrial protein extracts were prepared by solubilizing heart mitochondria in the same RIPA buffer. For electrophoresis, proteins were reduced and denatured in 1X Laemmli buffer, resolved on 10% SDS-PAGE gels, transferred to PVDF membranes, immunodetected with antibodies, and imaged using a ChemiDoc system (BioRad). Primary antibodies used in the study were: anti-acetylated lysine (PTM Biolabs, 1:1000, and Cell Signaling Technology #9441, 1:1000), anti-malonylated lysine (PTM Biolabs, 1:1000), anti-succinylated lysine (PTM Biolabs 1:1000), anti-glutarylated lysine (PTM Biolabs, 1:1000), anti-pAMPK (Cell Signaling Technology #2532S 1:1000), anti-AMPK (Cell Signaling Technology #2535S, 1:1000) anti-SIRT5 (Cell Signaling Technology #8782, 1:1000), , anti-IDH2 (Proteintech), anti-Porin (Abcam, ab14734, 1:1000), anti-α-tubulin (Proteintech 11224-1-AP, 1:1000) and anti-GAPDH (Fitzgerald, 10R-G109A, 1:10000).

### Immunoprecipitation

Mouse heart tissue was solubilized in immunoprecipitation buffer [20 mM Tris-HCl (pH 7.5) 250 mM NaCl, 1% Triton X-100, Nicotinamide 2.4mg/ml, 0.5 mM dithiothreitol, protease inhibitor cocktail (Roche)]. To assess IDH acetylation or malonylation, 1mg of total heart proteins were immunoprecipitated with the following antibodies: anti-IDH2 (Bethyl Laboratories) or 3.5ug rabbit IgG (Bethyl Laboratories) for 3 h at 4°C, followed by an overnight incubation with protein A-agarose beads (Santa Cruz) at 4°C. Immunoprecipitated proteins were resolved by acrylamide gel electrophoresis and immunoblotted with antibodies against either acetylated lysine (Cell Signaling) or malonylated lysine (Cell Signaling). VeriBlot (Abcam, 1:1000) was used as an IP Detection Reagent, blots were visualized by chemiluminescence using the SuperSignal West Femto Maximum Sensitivity Substrate (Thermo scientific) and imaged using a ChemiDoc system (BioRad). To assess SIRT3 acetylation, the same immunoprecipitation protocol as described above was followed, except anti-acetylated lysine (Cell Signaling) or control rabbit polyclonal anti-HA (Proteintech) antibodies were used. Immunoblotting was conducted with anti-SIRT3 (Proteintech, 1:1000). A mouse anti-rabbit conformation specific secondary antibody (Cell Signaling) was used.

### Real-time PCR

Total RNA was extracted from heart tissue using the RNeasy Fibrous Tissue Mini Kit (Qiagen) and cDNA was generated using the High Capacity cDNA Reverse Transcription Kit (Applied Biosystems). RT-PCR was performed on a 7500 Real-Time PCR System (Applied Biosystems) using the iTaq Universal SYBR Green Supermix (BioRad). The ΔΔC(t) method was used to quantify the fold change of the target genes. The primer sets used were: ATF4 (Forward, GCCGGTTTAAGTTGTGTGCT; Reverse, CTGGATTCGAGGAATGTGCT) (67), ATF5 (Forward, GGGTCATTTTAGCTCTGTGAGAGAA; Reverse, ATTTGTGCCCATAACCCCTAGA) (68), and RPS20 (Forward, AACAAGTCGGTCAGGAAGC; Reverse, TCCGCACAAACCTTCTCC).

### Amplex Red Assay

Isolated cardiac mitochondrial fractions were solubilized in MS-EGTA buffer, and protein concentrations were quantified by Bradford assay (BioRad). Measurements of hydrogen peroxide production was quantified using an Amplex Red assay (Invitrogen #A22188) according to manufacturer’s instructions.

### IDH2 Assay and Structural Analyses

Enriched mitochondrial fractions were prepared from HEK293 cell lines by differential centrifugation using cellular mitochondrial isolation buffer (250mM sucrose, 20mM HEPES, 10mM KCl, 1.5mM MgCl2, 1mM EDTA, 1mM EGTA, 1mM DTT, pH 7.5) supplemented with 2.4mg/mL nicotinamide and a combined phosphatase and protease inhibitor cocktail (Thermo Fisher) or cardiac mitochondria prepared as described above. Measurement of IDH activity on the enriched mitochondrial fractions was conducted using an IDH activity assay (Sigma Aldrich) according to the manufacturer’s instructions. Protein concentrations were quantified by Bradford assay, and IDH2 activity was normalized to the amount of mitochondrial protein used in the assay.

IDH2 (PDB ID: 5H3F (43)) structural data was retrieved from the RCSB Protein Data Bank (69) and all figures of structures were created with the Mol* software (70).

## ACKNOWLEDGEMENTS

Research reported in this publication was supported in part by the Emory Integrated Proteomics shared resource of Winship Cancer Institute of Emory University and NIH/NCI under award number P30CA138292. The content is solely the responsibility of the authors and does not necessarily represent the official views of the National Institutes of Health. The acyl-CoA measurements were performed by the Southeast Center for Integrated Metabolomics (SECIM) at the Sanford Burnham Prebys Medical Research Institute, Orlando, Florida.

---

**Figure S1.**
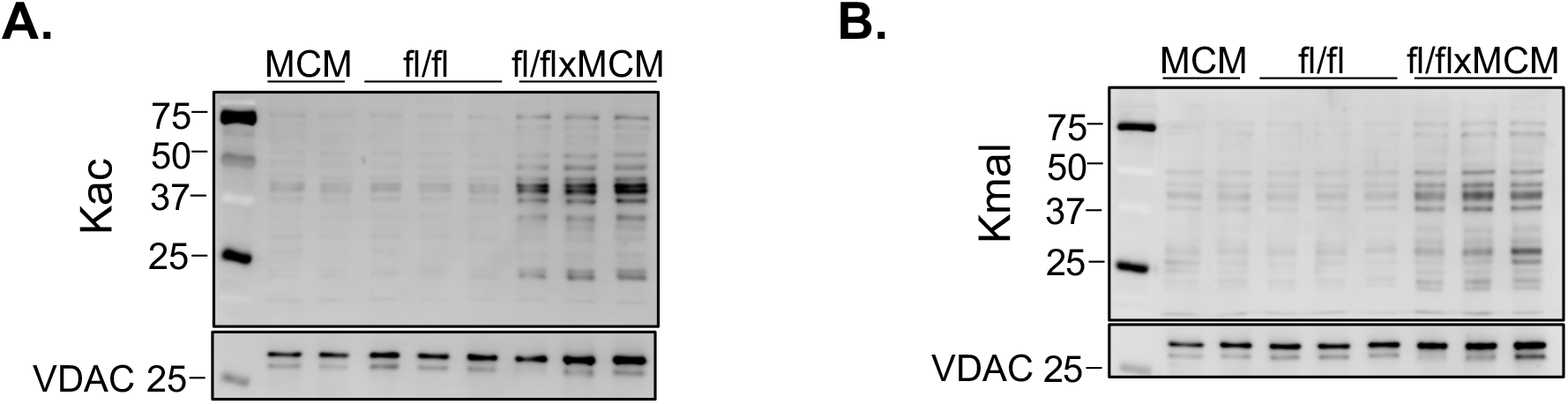
SLC25A3 deletion increases acetylation and malonylation of mitochondrial proteins. Western blot of (A) acetylated (Kac) and (B) malonylated (Kmal) lysine residues in cardiac mitochondria isolated from MCM, Slc25a3^fl/fl^ and Slc25a3^fl/flxMCM^ mice. VDAC was used as a mitochondrial protein loading control.

**Figure S2.**
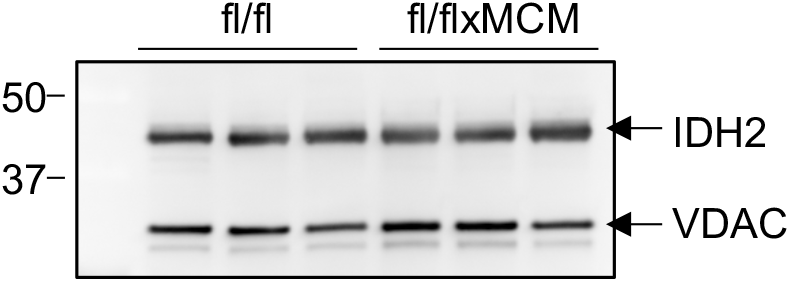
IDH2 expression is unchanged with SLC25A3 deletion. Western blot of IDH2 expression in isolated cardiac mitochondria from, Slc25a3^fl/fl^ and Slc25a3^fl/flxMCM^ mice 10 weeks following tamoxifen adminiWestern blot of IDH2 expression in isolated cardiac mitochondria from, Slc25a3^fl/fl^ and Slc25a3^fl/flxMCM^ mice 10 weeks following tamoxifen administration. VDAC was used as a mitochondrial protein loading control.

